# A framework for designing splice-junction experiments in deep 3’ single-cell RNA sequencing

**DOI:** 10.64898/2026.09.22.753522

**Authors:** Aabida Saferali, Congjian Liu, Andrea S Vendrame, Kamakshi Bankoti, Jose Santos Cabrera, Yohannes Tesfaigzi, Peter J. Castaldi

## Abstract

Alternative splicing is cell-type-specific and disease-relevant, but single-cell RNA sequencing is optimized for gene-level quantification, and how sequencing depth governs splice-junction recovery in the dominant 3’-biased chemistries has not been quantified, so experiments cannot be designed to a depth target and datasets cannot be interpreted against expected recovery. We generated deep single-cell RNA sequencing of human airway epithelial cells (12 samples, 102 to 803 million uniquely mapped reads per sample) and benchmarked junction recovery in five annotated samples against matched bulk RNA sequencing from a 190-donor cohort. Junction detection did not saturate with depth, and validity was governed by read support rather than annotation status. Recovering 90% of the observed full-depth junction complement required approximately 52,250 reads per cell, several-fold more than gene-level analysis. Quantifying splicing was more depth-limited than detecting it, and rare cell types were limited by cell number rather than depth. Against matched bulk, single-cell recall plateaued at a practical ceiling set by sample size and sparse sampling, and was independent of distance from the 3’ end. These results establish a framework for designing splicing experiments in deep 3’-biased single-cell RNA sequencing, and provide an airway-epithelial dataset with matched bulk for benchmarking splicing methods.

## Introduction

Alternative splicing generates functional diversity across nearly all human protein-coding genes and is regulated in a cell-type-specific manner [1, 2]. In the airway epithelium, splicing regulation contributes to cell identity, differentiation, and the response to environmental stressors [3]. In disease, altered splicing has been implicated in chronic obstructive pulmonary disease [4, 5], cystic fibrosis [6], and idiopathic pulmonary fibrosis [7, 8], often through cell-type-specific mechanisms that bulk analyses cannot resolve.

Single-cell RNA sequencing has transformed the study of cell-type-specific gene expression, but its application to splicing remains limited. Several tools for single-cell splicing quantification now exist [9–11], yet how much sequencing depth is required for reliable junction detection has not been quantified, so experiments cannot be designed to a depth target and existing datasets cannot be interpreted against an expected recovery.

Sequencing-depth requirements for gene-level quantification are relatively well established. Previous work has shown diminishing returns in gene-level information beyond approximately 15,000 reads per cell in droplet-based scRNA-seq, while current 10x Genomics guidelines recommend minimum sequencing depths of 20,000 read pairs per cell for standard 3’/5’ gene-expression libraries and 10,000 read pairs per cell for Flex [12–15]. In contrast, depth requirements for isoform-level analysis have primarily been characterized in full-length, plate-based protocols, where Westoby et al. found substantially improved isoform detection at approximately 4 million compared with 1 million reads per cell, with incomplete detection persisting even at the higher depth [16]. Between these regimes lies the depth range commonly used in droplet- and combinatorial-based single-cell studies, yet the relationship between sequencing depth and splice-junction recovery within this range remains poorly characterized.

The dominant high-throughput single-cell chemistries also impose constraints on transcript coverage. Conventional 10x Genomics 3’ and combinatorial indexing methods generate coverage strongly enriched toward transcript 3’ ends, whereas 10x Flex uses predefined transcript-specific probe pairs and therefore samples targeted regions of each gene [14]. Sparse and nonuniform transcript coverage can introduce coverage-dependent biases into estimates of alternative splicing [17]. Nevertheless, computational approaches such as SICILIAN have demonstrated that biologically meaningful splice-junction detection is achievable from 3’-biased single-cell data [10], leaving open how deep is deep enough for a given experimental goal.

Existing resources are not designed to define sequencing-depth requirements for splice-junction recovery. Public single-cell atlases are typically sequenced at depths far below those at which isoform recovery continues to improve, while existing studies of splicing in droplet-based data have focused on junction-detection accuracy at a fixed sequencing depth rather than systematically varying depth [10]. Conversely, studies with sufficient transcript coverage for detailed isoform analysis have largely relied on lower-throughput, full-length methods such as Smart-seq2 and Smart-seq3 [16, 18, 19]. Thus, no existing study has systematically evaluated how sequencing depth affects splice-junction recovery across the depth range relevant to contemporary high-throughput single-cell experiments.

Here we generated such a dataset in human airway epithelial cells differentiated at the air-liquid interface, using the ScaleBio combinatorial chemistry, with 12 samples sequenced deeply across a range of depths and matched bulk RNA sequencing from the same donors. We use this dataset to quantify how sequencing depth governs splice-junction recovery in single-cell RNA sequencing and to characterize the trade-offs that determine effective experimental design.

From these analyses we establish four elements of an experimental-design framework for splicing analysis in deep 3’-biased single-cell RNA sequencing: (1) depth targeting, matching sequencing depth to whether the analytical goal is junction detection or splicing quantification; (2) budget allocation, trading reads per cell against cell number as a function of cell-type abundance; (3) practical constraints, quantifying the recovery ceiling that more single-cell sequencing alone does not cross; and (4) chemistry considerations, testing whether 3’-bias produces a position-dependent blind spot for splicing analysis.

Beyond the framework, we show that the resulting data recover biologically coherent, cell-type-specific splicing, and we release a deeply sequenced human airway epithelial single-cell dataset as a resource for benchmarking future methods.

## Methods

### Study samples and single-cell RNA sequencing

Single-cell profiling was performed on primary human airway epithelial cells from the HAEC-185 cohort. Briefly, airway epithelial cells were collected from the large airways of 190 individuals over multiple years during studies of the regulators of mucus cell hyperplasia, and were isolated from human lung tissue obtained primarily from organ donor lungs deemed unsuitable for transplantation. All subjects or their authorized representatives provided written informed consent, and the use of human airway epithelial cells was approved by the Institutional Review Board at Brigham and Women’s Hospital. Airway basal cells are the progenitor cells that maintain and repair the airway epithelium. Matched bulk RNA sequencing of basal airway epithelial cells was available for the same cohort and was used as an orthogonal reference (below).

For single-cell profiling, basal airway epithelial cells from a subset of donors were maintained in culture and driven to differentiation under air-liquid interface conditions as previously described [20], and the differentiated cells were profiled by single-cell RNA sequencing using the ScaleBio ScaleRNA assay. A total of 12 samples were sequenced. Cell-type annotation and a matched bulk donor were available for five of these, the untreated primary samples, which were therefore the samples carried into the depth, cell-type, and recall analyses; all 12 contributed to the sample-level junction-yield and quality summary (Fig 2). Reads were aligned to the GRCh38 reference and quantified with STARsolo (STAR v2.7.10b) [21], using a 29-base cell barcode and an 8-base unique molecular identifier (UMI), forward strand orientation, and the GeneFull_Ex50pAS feature to retain both exonic and intronic reads. STARsolo alignments were written with corrected cell-barcode (CB) and UMI (UB) tags, which were used for all downstream cell-level analyses. For the recall analyses, five single-cell samples were matched to their corresponding bulk donor by a sample-to-donor crosswalk, enabling a within-donor comparison.

### Quality control and doublet removal

Genes detected in fewer than three cells were removed, and cells were retained on the basis of per-sample quality thresholds. The number of detected genes was required to exceed a lower bound defined as the median minus three median absolute deviations (MADs) on the log scale, a minimum library complexity was imposed, and the mitochondrial read fraction was required to fall below a per-sample upper threshold, defined as the median plus three MADs and capped at 25%. A fixed mitochondrial cutoff was deliberately avoided, because the mitochondrial read fraction is elevated in metabolically active ciliated cells; an adaptive per-sample threshold prevents the biased removal of these cells. In these differentiated airway cultures the mitochondrial fraction was low (per-sample median around 1%), so the adaptive value, rather than the 25% cap, set the threshold in every sample. Doublets were identified and removed within each sample using scDblFinder [22]. Cell-type annotation was performed on the five untreated primary samples, yielding 23,679 high-quality single cells.

### Normalization, integration, and clustering

Counts were normalized and variance-stabilized with SCTransform [23] using the glmGamPoi method [24]. Principal component analysis was performed on the corrected expression values, and the principal components were integrated across samples using Harmony [25] with the sample of origin as the batch covariate. Clustering was performed on the Harmony embedding using the Louvain algorithm. The clustering resolution was selected objectively rather than by inspection: across a grid of resolutions from 0 to 3 in steps of 0.05, the mean silhouette width of the resulting clusters was computed on the Harmony embedding, and a bootstrap confidence interval was obtained by resampling cells (50 replicates at 80% subsampling). The resolution maximizing the bootstrapped mean silhouette width was selected. Cluster stability was assessed by a clusterboot-style Jaccard analysis (30 bootstrap replicates), and clusters with a mean Jaccard coefficient of at least 0.75 were considered stable.

### Cell type annotation

Cell types were assigned by combining marker-based scoring with reference mapping. First, each cluster was assigned to an epithelial lineage using module scores computed over curated marker sets for basal, club, goblet, ciliated and deuterosomal, and rare (ionocyte, tuft, and pulmonary neuroendocrine) cell types. Second, cells were mapped onto a reference restricted to airway epithelium: the Human Lung Cell Atlas (HLCA) Azimuth reference [26] was subset to epithelial cells after excluding alveolar cell types, and query cells were projected onto the reference by anchor-based transfer of the finest-level HLCA labels. We restricted the reference to airway epithelium because mapping onto the full lung reference produced implausible assignments, including several thousand spurious ionocytes. Ambiguous or heterogeneous clusters were refined by iterative subclustering, with each candidate subcluster accepted only if it met predefined gates (Jaccard stability of at least 0.75, at least 50 cells, at least eight differentially expressed genes, and no single donor contributing more than 70% of its cells, so that donor-specific effects were not mistaken for a distinct cell state). Annotation yielded 23,679 cells spanning seven epithelial cell types (basal, club, ciliated, goblet, squamous, deuterosomal, and Transitional). Barcode-to-cell-type assignments were matched to the STARsolo CB tags for the junction-level analyses. All single-cell processing was performed in R 4.6 with Seurat 5.5 [27] together with the SCTransform/glmGamPoi, Harmony, scDblFinder, bluster, and Azimuth workflows described above.

### Splice junction detection

Two junction sources were used. The per-sample yield and composition summaries (Fig 2), reported across all 12 samples, were taken directly from STAR’s SJ.out.tab output; the depth, recall, and detectability analyses used a custom cell-barcode-aware extractor, run on the five annotated samples, that recovers the per-cell and per-cell-type junction support SJ.out.tab does not provide.

For the extractor, a spliced read was defined as any read containing an N operation in its CIGAR string, with at least eight matched bases immediately flanking each gap, an intron length between 50 and 500,000 bp, and a cell barcode present in the cell-type annotation; the corresponding intron coordinates were computed in one-based coordinates consistent with the STAR SJ.out.tab convention. Each read was assigned to a cell type using its CB tag, and junction read support was counted both as the number of uniquely mapped reads and as the number of UMI-deduplicated molecules (distinct CB and UB combinations). Junctions were quantified per cell type and in aggregate (pseudobulk). To validate the extraction, we compared the resulting junctions to the STAR SJ.out.tab calls for the same sample and found 92.5% concordance for junctions supported by at least five reads and 97.1% for junctions supported by at least ten reads; because the extractor and SJ.out.tab apply different filters, this figure reflects overall agreement between the two pipelines rather than the parser in isolation. Junctions were classified as annotated or novel using the annotation flag from STAR SJ.out.tab. Detected junctions were assigned to genes by containment within GENCODE (v49) gene models [28], and the number of genes contributing at least one junction supported by at least five reads was tabulated per sample.

### Depth downsampling and saturation analysis

To characterize how junction detection depends on sequencing depth, we downsampled the spliced reads to a range of fractions (5% to 100%) and recounted the junctions detected at each depth, separately for annotated and novel junctions. Downsampling was performed at the read level and was UMI-aware, so that a junction was considered detected when at least one UMI-supported read survived. Sequencing depth was expressed as the number of uniquely mapped reads, obtained from the STAR Log.final.out file for each sample, and depth per cell was obtained by dividing this number by the count of annotated cells in the sample.

### Reads-per-cell versus cell-number analysis

To distinguish the contributions of sequencing depth and cell number to junction recovery, we subsampled cells and reads independently. For a grid of cell fractions and read fractions, we sampled the corresponding number of cells, downsampled their reads to the target fraction, and counted the junctions detected. Reads per cell were computed as the product of the read fraction and the total uniquely mapped reads divided by the number of cells. Recovery was evaluated both across the full grid and along lines of constant total sequencing budget.

### Bulk RNA sequencing and junction recall

Matched bulk RNA sequencing was available for 190 basal airway epithelial donors from the HAEC-185 cohort, sequenced to a median of 35.5 million read pairs per donor and aligned with STAR as part of an nf-core/rnaseq workflow [29]. A bulk junction catalog was constructed by taking the union of the per-donor STAR SJ.out.tab junctions, and each junction was annotated with the number of donors in which it was detected (with at least one uniquely mapped read) and its total read support. Because both datasets were aligned with STAR, junctions were compared directly, after removing the chromosome-name prefix from the bulk coordinates. Single-cell recall was defined as the fraction of bulk junctions recovered by the single-cell data, and was evaluated as a function of single-cell sequencing depth and of the bulk detection frequency (the number of donors supporting each junction).

### Distance from the transcript 3’ end

To test whether junction recovery depended on position within the gene, each bulk junction was assigned to the containing GENCODE (v49) gene, and the distance from the junction to the gene 3’ terminus was computed. To account for isoform variation, the analysis was repeated using the distance to the nearest 3’ terminus among all transcripts containing the junction. Single-cell recall was then evaluated in bins of this distance, stratified by bulk detection frequency to control for expression.

### Gene body coverage

Gene body coverage was computed for the five annotated samples. Briefly, each protein-coding gene between 3 and 100 kb in length was divided into 100 bins from the 5’ to the 3’ end, in a strand-aware manner, and each read was assigned to a single bin by its reference midpoint (rather than contributing per-base coverage across the bins it spans). Per-gene profiles were normalized to sum to one so that each gene contributed equally, and averaged across all genes with at least 50 reads.

### Intron clustering and differential splicing across cell types

To test for cell-type-specific splicing, splice junctions were grouped into intron clusters following the LeafCutter definition [30] in which junctions that share a 5’ or 3’ splice site are connected into the same cluster. Clusters with at least two junctions and at least 20 UMI-deduplicated molecules were retained, and molecule counts were tabulated for each junction in each combination of donor and cell type. Differential junction usage across the six cell types present in all five donors (basal, Transitional, ciliated, club, goblet, and squamous) was tested with edgeR (v4.10.0) [31] using the diffSpliceDGE function, with the intron cluster as the grouping unit and the donor of origin included as a blocking covariate, so that cell types were compared against biological replicates rather than against individual cells. Before testing, junctions with fewer than 10 pseudobulk molecules summed across all donor-by-cell-type samples were removed, and each cluster was again required to retain at least two junctions after this filter. An omnibus test across cell types was formed for each cluster by taking the minimum of the cell-type Simes p-values across the cell-type coefficients, applying a Bonferroni correction for the number of coefficients, and controlling the false discovery rate across clusters with the Benjamini and Hochberg procedure. Clusters with an adjusted p-value below 0.05 were considered differentially spliced, and were assigned to genes by the most frequent GENCODE (v49) gene containing their junctions. Gene-set enrichment of the differentially spliced genes was assessed with fgsea (v1.38.0) [32] over the MSigDB Hallmark, Reactome, and Gene Ontology collections [33, 34], with genes ranked by their differential-splicing significance.

### Alternative-splicing quantification versus sequencing depth

To distinguish the depth required to detect a junction from the depth required to quantify splicing, UMI-deduplicated molecules were downsampled within each sample. Each molecule was assigned a stable uniform value and was retained at a given depth fraction when that value fell below the fraction, producing nested subsamples spanning 5% to 100% of depth. At each depth we counted the junctions detected (at least one molecule), the intron clusters that were quantifiable (at least two junctions each supported by at least ten molecules, so that a usage ratio could be estimated), and the accuracy of the percent-spliced-in (PSI) estimate. For each cluster, PSI was defined for its two most abundant junctions as the fraction contributed by the first, and its accuracy at each depth was summarized as the mean absolute deviation from the full-depth value, averaged over clusters quantifiable at full depth.

### Comparison with MARVEL

To assess whether the cell-type splicing signal was robust to the quantification method, junction usage was quantified independently with MARVEL (v2.0.5) [9]. A MARVEL single-cell object was constructed from the per-cell STARsolo splice-junction count matrices together with the gene expression matrices and cell-type annotations from the Seurat object, and splice junctions were annotated against the GENCODE (v49) annotation. After junction annotation and validation, differential junction usage between ciliated and basal cells was tested with the MARVEL permutation procedure (100 iterations). Concordance with the intron-cluster quantification was assessed on the junctions quantified by both methods, by correlating the ciliated-minus-basal PSI change with the Spearman correlation and by measuring the agreement in the direction of change among junctions with an absolute PSI change of at least 0.05. MARVEL defines junction usage relative to total gene expression, whereas the intron-cluster approach defines it relative to the competing junctions within a cluster.

### Annotation-guided re-alignment

To test whether missed bulk junctions reflect genuine under-sampling or a failure to align reads that were present, the cDNA reads of the five annotated samples were re-aligned with STAR twice: once against the standard genome index, and once with the confident bulk junctions (those detected in at least five donors) inserted into the splice-junction database on the fly. STAR aligns reads across an annotated junction even with a short overhang that would otherwise be soft-clipped, so a difference between the two alignments isolates the junctions recoverable by annotation-guided alignment. Recall of the confident junctions (detected with at least five uniquely mapped reads) was compared between the two alignments.

### Detectability model

To predict junction detection from experimental parameters, we assembled a training grid by subsampling, within each cell type, both the cells (retaining a stable uniform fraction) and the UMI-deduplicated molecules (nested read-level downsampling), and recording, for each junction expression bin, the fraction of reproducible bulk junctions (present in at least two donors) that were detected. Junction expression was taken as the mean bulk read support per donor, cell number as the retained number of cells of the type, and reads per cell as the read fraction multiplied by the pooled per-cell depth. Because a junction is detected when at least one of its molecules is captured, Poisson sampling implies that the detection probability is a complementary-log-log function of the logarithms of expression, cell number, and reads per cell; we fitted this as a binomial generalized linear model with a complementary-log-log link (R 4.6), weighting each condition by its number of bulk junctions. Generalization was assessed by leave-one-cell-type-out cross-validation, in which the model was trained on all cell types but one and used to predict the held-out cell type, and agreement between predicted and observed detection was summarized as a count-weighted coefficient of determination and mean absolute error.

### Code and data availability

The single-cell RNA-sequencing data generated in this study have been deposited in the Gene Expression Omnibus (GEO) under accession GSE345128 and will be made publicly available upon publication. The deposit includes raw FASTQ files and STARsolo count matrices for the five untreated primary samples used for the depth, cell-type, and recall analyses. The bulk RNA-sequencing data are part of an ongoing companion study and will be deposited in a public repository following publication of that study. To enable reproduction of the analyses reported here, processed splice-junction data derived from the bulk RNA-sequencing dataset, including the junction counts used for comparison with the single-cell data, will be made publicly available with this article. Analysis code will be made publicly available on github.

## Results

### A deep, cell-type-annotated single-cell atlas of the airway epithelium

We generated single-cell RNA sequencing data from differentiated human airway epithelial cells of the HAEC-185 cohort using the ScaleBio ScaleRNA assay, sequencing 12 samples to a median of 438 million uniquely mapped reads per sample (range 102 million to 803 million; 5.10 billion in total), far above the depth used for routine gene-level profiling. To establish the cellular landscape underlying the splicing analyses, we processed the five untreated, non-transformed samples through quality control, doublet removal, normalization, integration, and annotation, yielding 23,679 high-quality single cells at a median of 58,056 uniquely mapped reads per cell (range 32,274 to 133,068). Clustering and reference-guided annotation resolved seven epithelial cell types (basal, club, ciliated, goblet, squamous, deuterosomal, and Transitional), spanning the expected common-to-rare gradient from abundant basal and club cells to the small Transitional population and rarer deuterosomal cells (Fig 1).

**Figure 1.**
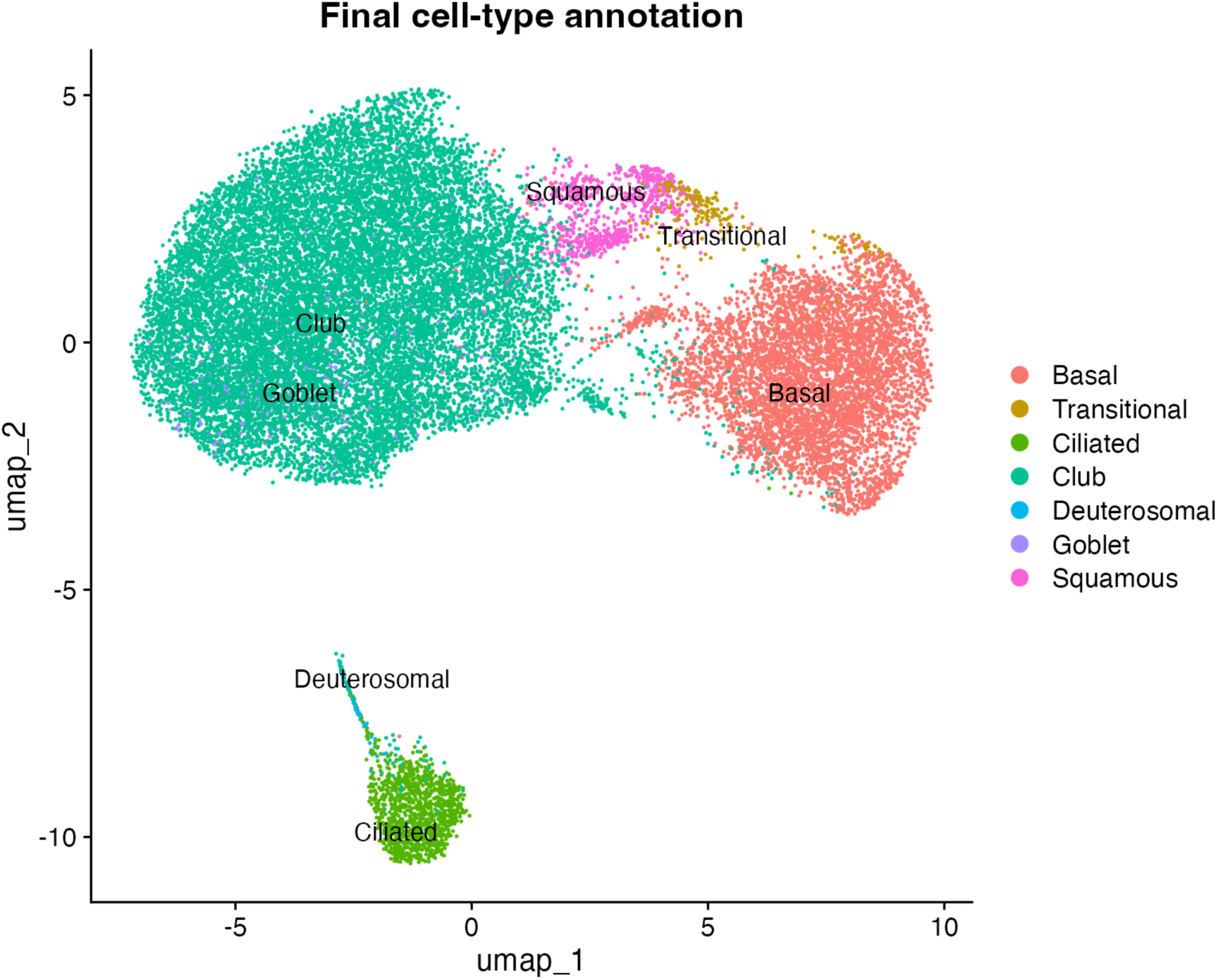
Cell-type annotation of differentiated airway epithelial cells. UMAP embedding of 23,679 high-quality single cells from five samples, colored by the seven annotated epithelial cell types (basal, Transitional, ciliated, club, deuterosomal, goblet, and squamous). These cell types define the common-to-rare gradient used in the depth analyses.

### Deep 3’ single-cell data yields abundant, high-confidence splice junctions

Deep 3’ single-cell RNA sequencing recovers abundant splice junctions (Fig 2). Extracting junctions directly from the STARsolo alignments, we found a median of 121,590 junctions per sample at a five-read threshold (range 69,126 to 151,681), arising from a median of 14,662 genes. A median of 90.8% used a canonical splice motif and 77.0% were annotated in the reference; because the airway epithelium is comparatively understudied at the transcriptome level, we treat annotation status as a description of novelty rather than a measure of validity, and benchmark that novelty against the matched bulk catalog below. As expected for sparse single-cell data, the read-support distribution was heavy-tailed, with 29.3% of junctions supported by only a single read. Deep 3’ single-cell RNA sequencing therefore yields abundant and largely high-confidence junctions; the challenge is how much sequencing is needed to sample them completely.

**Figure 2.**
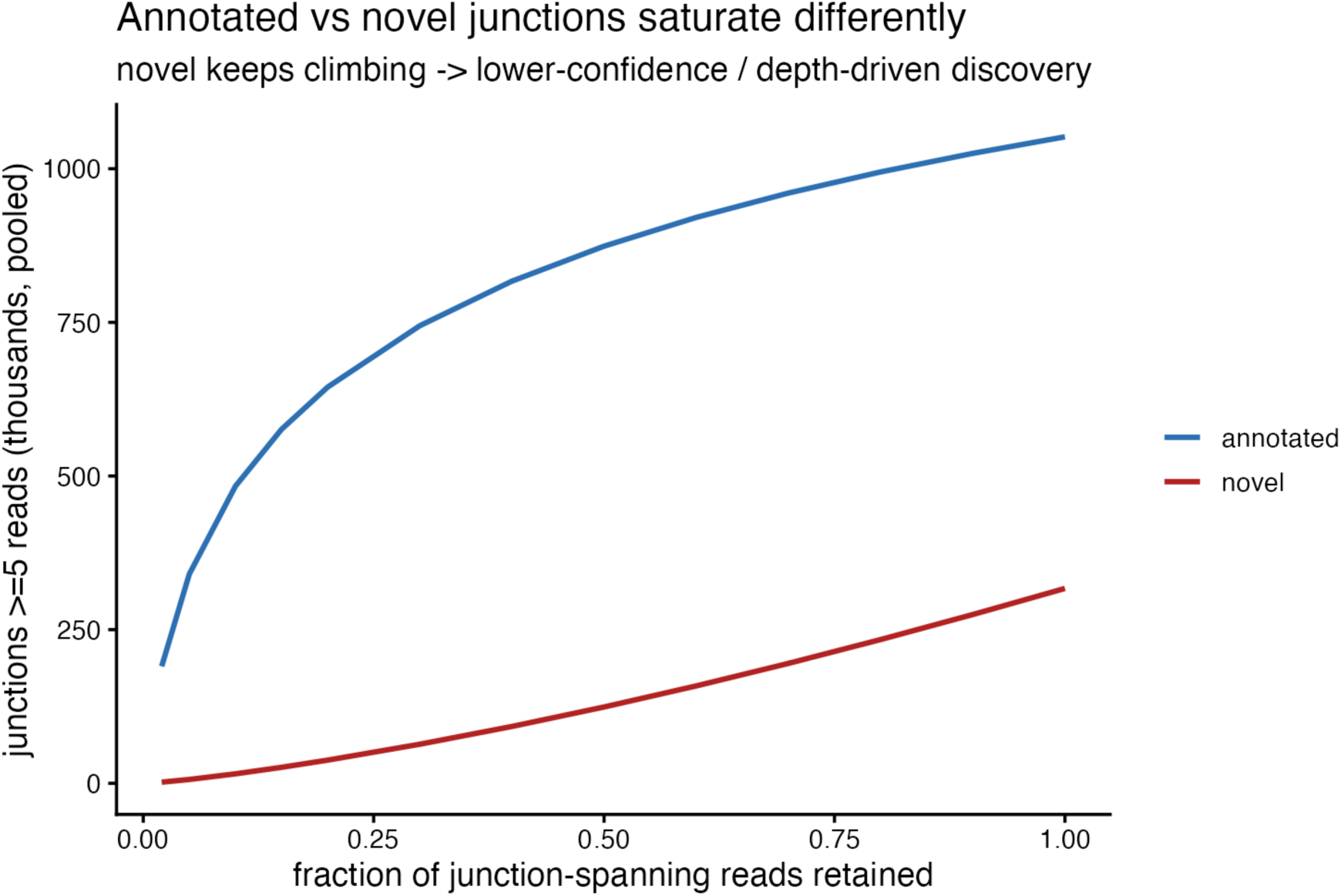
Splice-junction yield and quality in deep 3’ single-cell data. Per-sample counts of splice junctions supported by at least five reads, partitioned into annotated and novel. Across the 12 samples a median of 121,590 junctions were detected per sample, of which a median of 77.0% were annotated and 90.8% used a canonical splice motif.

### Junction detection does not saturate at the sequencing depths achieved

To quantify how junction detection depends on sequencing depth, we performed UMI-aware read-level downsampling within each sample and recounted the junctions detected at each depth (Fig 3). At full depth, samples yielded a median of 491,049 junctions (range 262,017 to 698,480), and total junction detection did not saturate across the depths we achieved: halving the sequencing depth still recovered 67% of the full-depth junctions, and even the deepest samples were still gaining junctions. This aggregate non-saturation is driven by the lowly expressed junctions: when detection is stratified by junction expression, the highly expressed junctions in a common cell type approach saturation (in club cells, detection of junctions expressed at 200 or more reads per donor reaches 85% and gains only about 5% relative over the final doubling of depth), whereas the lowly expressed junctions are far from saturated (reaching only about 6% detection at full depth, a 52% relative gain over the same doubling). Separating annotated from novel junctions showed that this additional depth is spent unevenly: between 10% and 100% of depth, annotated junctions grew 1.6-fold whereas novel junctions grew 6.1-fold.

**Figure 3.**
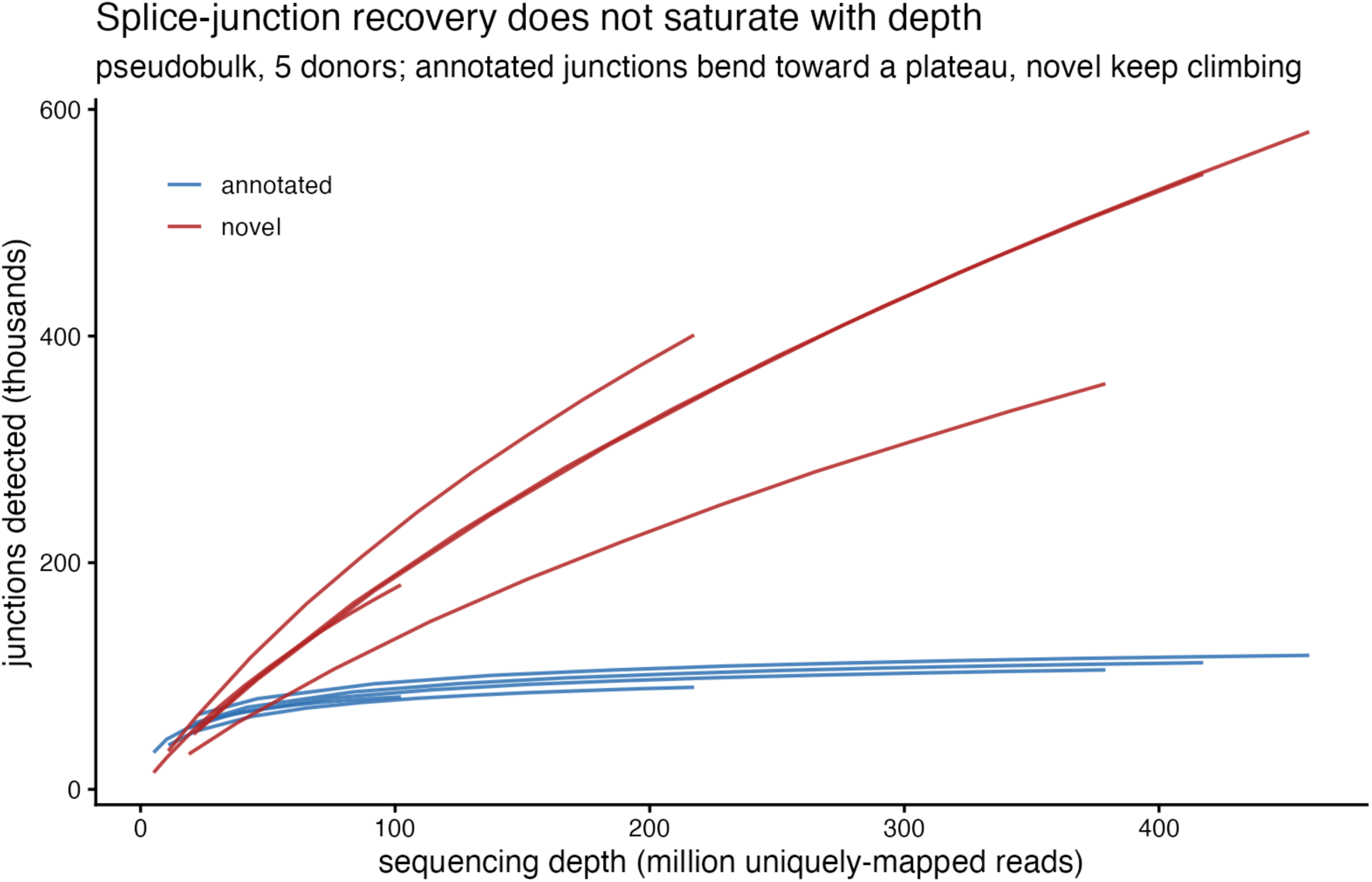
Junction detection does not saturate at the sequencing depths achieved. Within-sample UMI-aware read downsampling. Junctions detected as a function of sequencing depth, shown separately for annotated and novel junctions. Between 10% and 100% of depth, annotated junctions grew 1.6-fold whereas novel junctions grew 6.1-fold.

Rather than treat this novel growth as low-confidence noise, we asked what actually predicts whether a junction is real by benchmarking each single-cell junction against the matched bulk catalog (Fig S1). Read support, not annotation status, was the primary determinant of validity: the fraction of junctions confirmed in bulk rose monotonically with read support for both classes, from 10% for novel single-read junctions to 83% for novel junctions with at least 200 reads, while annotated junctions were confirmed at 83% even at a single read and at essentially 100% with deeper support.

A substantial and increasing fraction of the novel junctions are therefore real but simply lower expressed and absent from the reference annotation, and the genuinely uncertain population is the single-read calls (29.3% of detected junctions) rather than novel junctions as a class. Deep sequencing thus recovers a minority of genuine, lower-expressed junctions alongside a large tail of low-support calls: of the junctions gained between 10% and 100% of depth, only about 14% were confirmed in the matched bulk catalog (present in at least two bulk donors, pooled across the five annotated samples), reflecting their low read support rather than their novelty. Some of these low-support calls may additionally arise from regions that are intrinsically difficult to sequence or align rather than from low expression alone, a distinction that read-level evidence cannot fully resolve. Confidence is therefore best gated on read support rather than on annotation status.

### Junction recovery requires several-fold more depth than gene expression

Because both sequencing depth and cell number contribute to junction recovery, we subsampled cells and reads independently across a grid and measured junction recovery as a function of reads per cell (Fig 4A). Reaching 90% of the full-depth junction complement required approximately 52,250 reads per cell (median across the five donors, which individually ranged from 29,046 to 119,761), threefold to fourfold higher than the ∼15,000 reads per cell beyond which previous work observed diminishing returns in gene-level information. This higher requirement reflects that junction-level counts sit one to three orders of magnitude below gene-level counts (Fig S2), because a gene’s reads are distributed along its length and split among its junctions.

**Figure 4.**
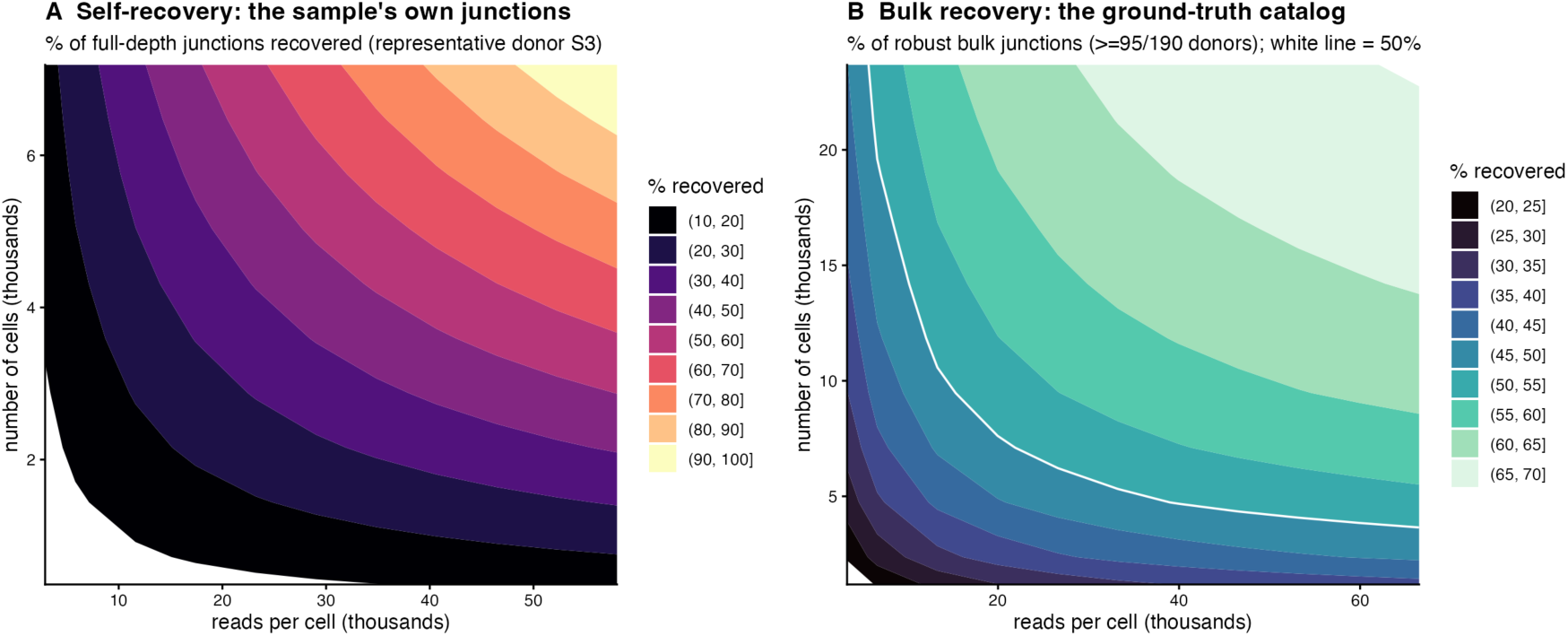
Reads-per-cell versus cell-number trade-off for junction recovery. Junction recovery as a joint function of reads per cell and number of cells, with cells and reads subsampled independently. (A) Recovery of the sample’s own full-depth junctions (representative donor S3); reaching 90% of the full-depth complement required approximately 52,250 reads per cell, several-fold higher than the approximately 15,000 reads per cell beyond which previous work observed diminishing returns in gene-level information. (B) Recovery of the ground-truth matched-bulk catalog (junctions in at least 95 of 190 donors, n = 194,492), pooled across the experiment; the white line marks 50% recovery. Both panels show the same reads-per-cell versus cell-number tradeoff.

Anchoring the same axes to the matched bulk turned this into concrete recovery targets (Fig 4B). Recovering 50% of the robustly expressed bulk junctions (those present in at least 95 of 190 donors; n = 194,492) required approximately 6,700 reads per cell when all 23,679 cells were used, rising to approximately 13,300 reads per cell at roughly 11,800 cells and approximately 39,900 reads per cell at roughly 4,700 cells; 50% recovery was unattainable below approximately 4,700 cells at any depth. Reported at a fixed depth with all 23,679 cells used, recovery of this robust catalog rose from 62% at 20,000 reads per cell to 69% at 50,000 and 70% at full depth, so panel B occupies the 60 to 70 percent band rather than approaching 100%: even the robustly expressed bulk junctions are not all recovered in single-cell data. Recovery of the broader catalog (junctions in at least 20 donors) plateaued at 47.4%, the practical ceiling characterized below.

### Quantifying splicing requires more depth than detecting a junction

Detecting a splice junction is less demanding than quantifying its usage. We therefore compared depth requirements for junction detection and alternative-splicing quantification by downsampling UMI-deduplicated molecules (Fig S3). Quantification lagged detection at every depth. At half of full depth (each sample downsampled to 50% of its reads), 78% of full-depth junctions were still detected, but only 68% of quantifiable alternative-splicing events were recovered, and the gap widened toward low depth (at 5% of depth, 32% of junctions but only 9% of quantifiable events). The precision of the PSI estimate, measured as the mean absolute deviation from each sample’s own full-depth value, improved only slowly with depth (0.037 at half depth, approaching zero only at full depth). Full-depth sequencing yielded a median of 6,564 quantifiable alternative-splicing clusters per sample. The depth required to measure splicing is therefore substantially greater than the depth required to catalog junctions, an important distinction for studies that aim to quantify, and not merely detect, alternative splicing.

Because that deviation measures precision against each sample’s own full-depth estimate rather than accuracy against an independent reference, we separately compared single-cell PSI to the mean PSI of the same intron clusters across the 190 bulk donors (Fig S4). For well-expressed clusters, single-cell PSI differed from the bulk mean by about 0.15 on average, and this difference barely narrowed with depth (0.153 at half depth to 0.151 at full depth for the most highly expressed clusters), indicating that the single-cell-versus-bulk PSI gap is dominated by the biological difference between the differentiated single-cell cultures and the basal bulk reference rather than by single-cell under-sampling. Across-donor PSI variability in single-cell data was nonetheless comparable to bulk for these clusters (standard deviation 0.037 across the five single-cell donors versus 0.031 across the 190 bulk donors). Depth therefore governs the precision of a single-cell PSI estimate, whereas its agreement with an independent bulk reference is set by biology.

### Recovery in rare cell types is limited by cell number, not depth

Applying the depth analysis across the common-to-rare cell-type gradient defined in Fig 1 showed that junction recovery tracks the number of cells available to aggregate (Fig 5). At full depth, the common club-cell population yielded 375,961 junctions, whereas the small Transitional population yielded only 16,700 junctions, a more than twentyfold difference driven by cell number rather than by sequencing depth. Because rare populations contribute few cells to pseudobulk aggregation, the absolute junction complement recoverable from these populations remains much smaller even at deep sequencing, indicating that adding reads cannot substitute for adding cells when the population of interest is small. Defining a separate junction universe for each cell type (its own full-depth complement) showed that this is a matter of scale rather than of differing saturation behavior: the fractional saturation curve was nearly identical across cell types, with 69% to 72% of each cell type’s complement recovered at half depth from the common club population to the rare deuterosomal population (Fig S5). The depth needed to reach a given fraction of a cell type’s junctions is therefore cell-type-independent, while the absolute number of junctions attainable, from 375,961 in club to 12,671 in deuterosomal, is set by cell number.

**Figure 5.**
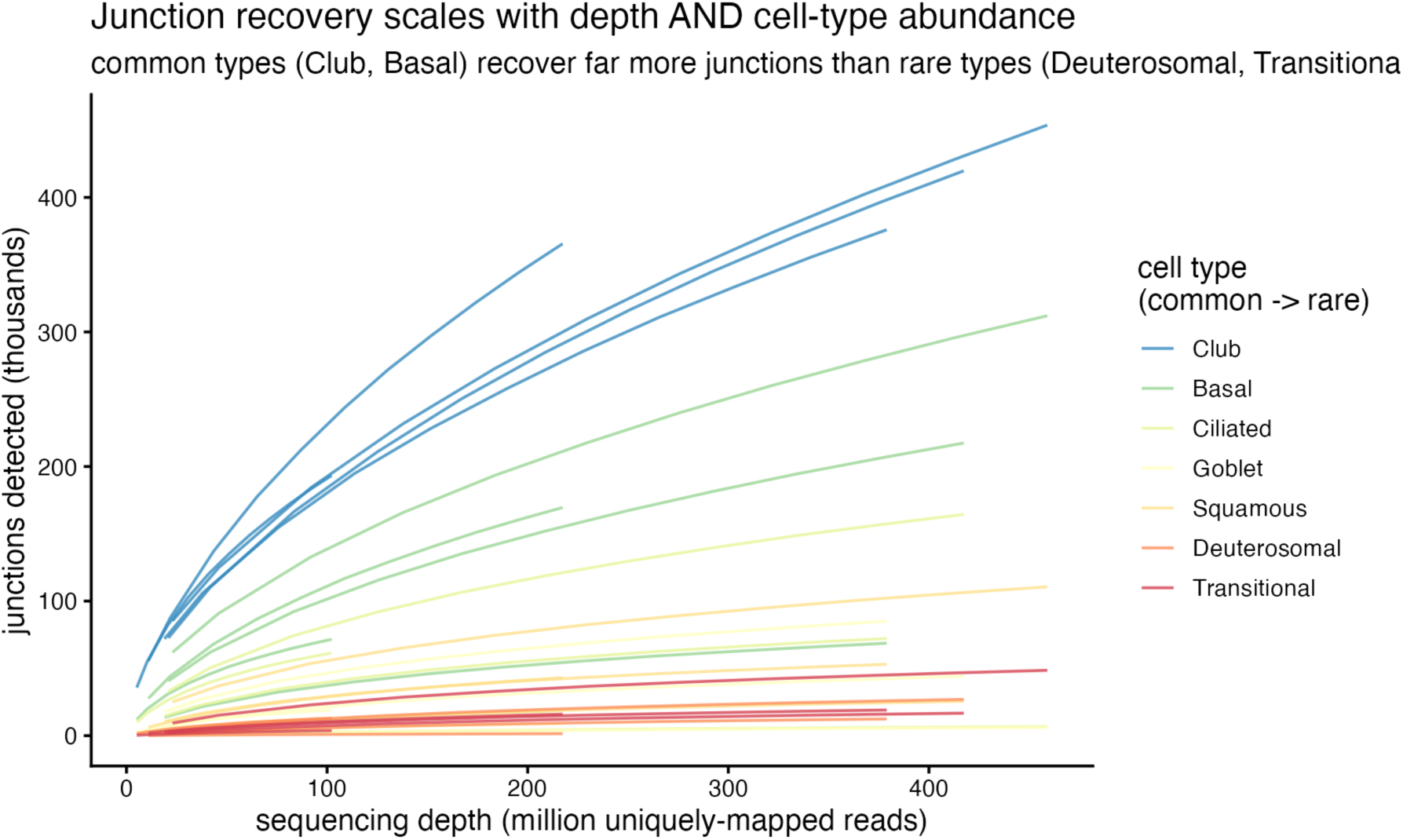
Junction recovery across the common-to-rare cell-type gradient. Junctions recovered per cell type as a function of depth. Recovery tracks the number of cells available for aggregation, from 375,961 junctions in the common club population to 16,700 in the small Transitional population, and rare populations remain far from saturation even at deep sequencing.

Detectability was jointly governed by cell-type abundance and junction expression level (Fig 6). Stratifying the reproducible bulk junctions by their expression, common cell types detected even lowly expressed junctions, whereas rare cell types recovered only the most highly expressed ones: club cells (15,363 cells) recovered 85% of the most highly expressed junctions (the same club-cell near-saturation reported in Fig 3) but only 15% of lowly expressed ones, and the rarest population (deuterosomal, 68 cells) recovered just 29% of even the most highly expressed junctions. A junction’s detectability in single-cell data can therefore be read directly from how abundant its cell type is and how highly it is expressed, providing a practical lookup for whether a given splicing event is recoverable in a planned experiment.

**Figure 6.**
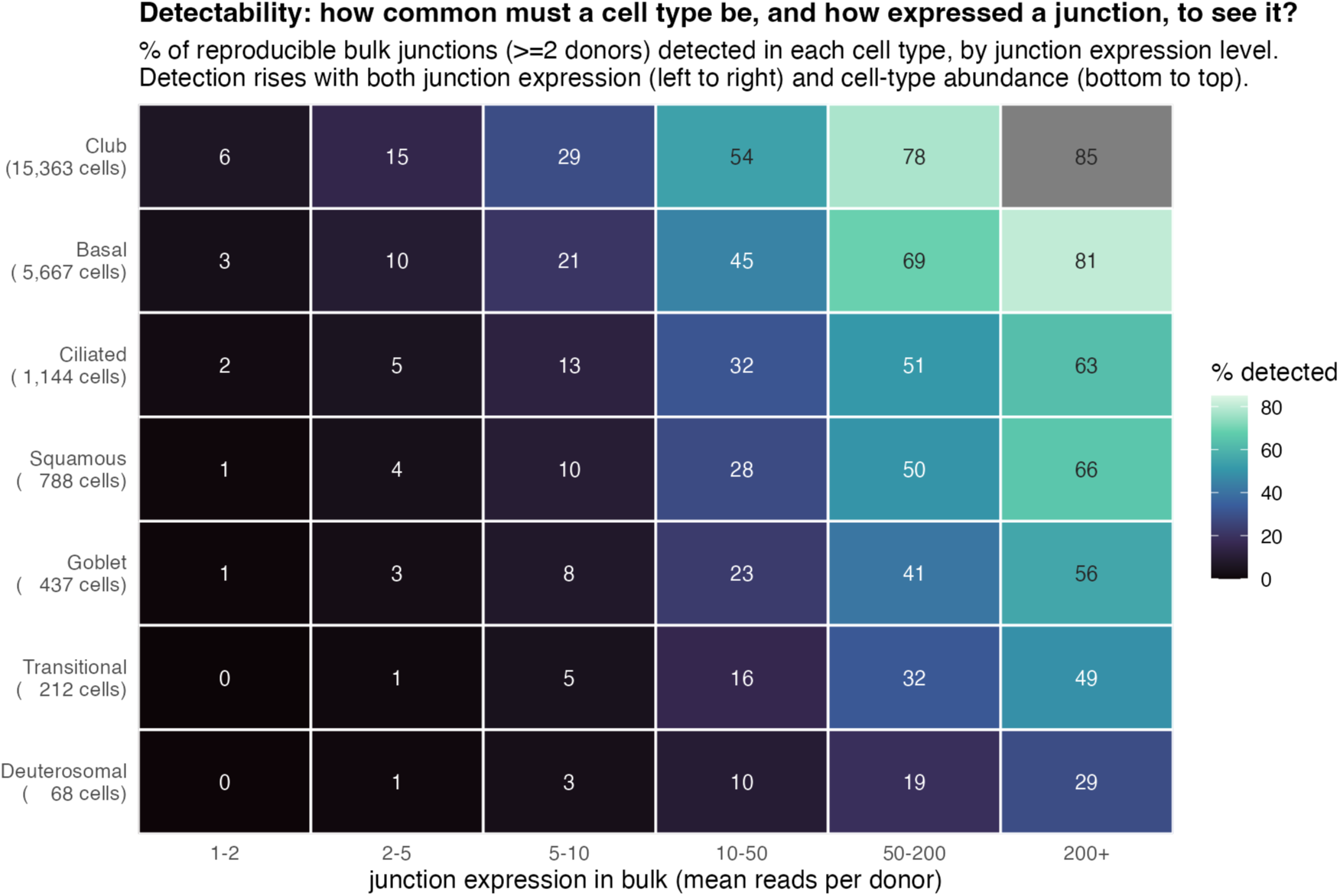
Detectability as a function of cell-type abundance and junction expression. Percent of reproducible bulk junctions (present in at least two donors) detected in each cell type, stratified by junction expression level (mean bulk reads per donor). Detection rises with both junction expression and cell-type abundance; cell types are ordered by the number of cells.

### Recall against matched bulk reveals a practical, not depth-limited, ceiling

To benchmark single-cell junctions against ground truth, we compared them to a catalog built from matched bulk RNA sequencing of 190 basal airway epithelial donors from the same cohort, sequenced to a median of 35.5 million read pairs per donor (Fig 7). Single-cell recall depended on the bulk detection-frequency threshold used to define the reference catalog, so we report it across a range of thresholds rather than as a single number. Recall of bulk junctions detected in at least 20 donors plateaued at 47.4% and was strongly stratified by how robustly a junction was expressed in bulk: recall rose from 47.4% for moderately supported junctions to 70.4% for junctions present in at least 95 donors and to 98.8% for universally detected junctions. Recall was higher for annotated than for novel junctions (47.4% versus 9.8%); because the bulk catalog is roughly 99% annotated, the novel comparison covers only about 1% of the catalog, and the gap is consistent with these novel bulk junctions being lower-expressed on average rather than less genuine. This plateau was not a depth artifact: doubling single-cell depth from 50% to 100% raised recall by only 4.0 percentage points, and a within-donor comparison of each single-cell sample against that same donor’s bulk reproduced it almost exactly (49.1%). The junctions that single-cell data miss are the lowly and variably expressed ones. Rather than an absolute limit, this is best read as a practical ceiling: the bulk reference draws on 38 times as many donors (190 versus 5) and a comparable total amount of sequencing (approximately 6.7 billion bulk reads), so its junction catalog is more exhaustive for reasons of sample size as much as chemistry. Treated as an exhaustive resource, the bulk data hold 807,447 junctions supported by at least five reads pooled across the 190 donors, of which the single-cell experiment recovered 311,053; closing the remaining gap would require more single-cell samples, not more reads, because within these five annotated samples ultra-deep sequencing was already performed.

**Figure 7.**
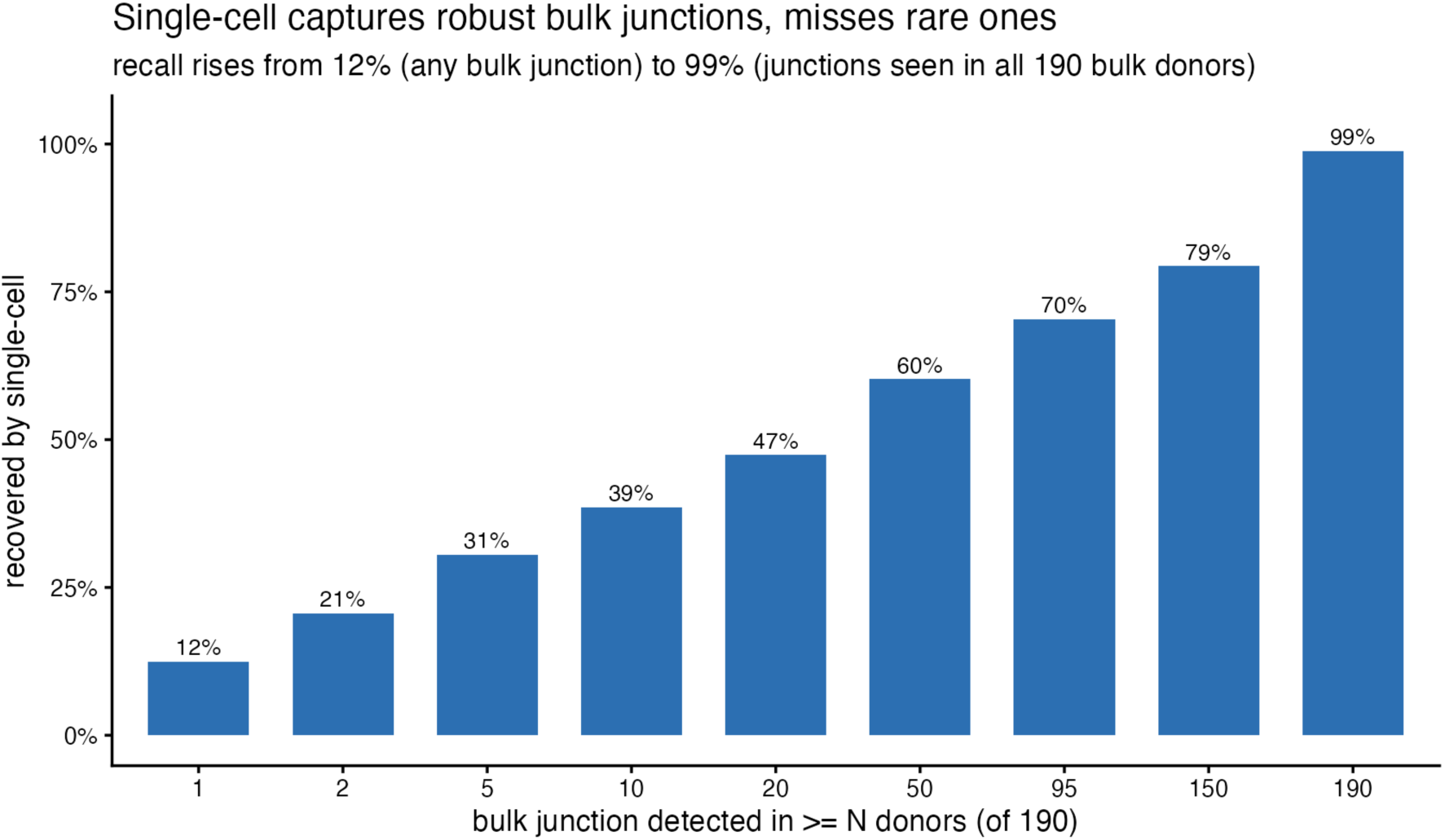
Recall of matched bulk junctions is set by expression robustness, not depth. Single-cell recall of bulk junctions stratified by the number of donors supporting each junction in the 190-donor bulk catalog. Recall rose from 47.4% for junctions in at least 20 donors to 98.8% for universally detected junctions, and was higher for annotated than novel junctions (47.4% versus 9.8%), reflecting the lower average expression of novel bulk junctions. Doubling single-cell depth added only 4.0 percentage points.

We further asked whether this ceiling reflects the genuine absence of the missing junctions rather than a failure to align reads that were present. Re-aligning the cDNA reads with the confident bulk junctions (those detected in at least five donors) supplied to the aligner as annotation, and comparing junction detection with and without this annotation, rescued only a small fraction of the missed junctions: recall of the confident set rose from 14.4% to 15.4% (a 6.9% relative increase), recovering between 3,424 and 9,464 junctions per sample. The great majority of missed junctions were therefore not recoverable by annotation-guided alignment, indicating that the recall ceiling reflects genuine under-sampling of junction-spanning reads rather than an alignment-sensitivity artifact.

### Junction recovery is governed by expression level, not distance from the 3’ end

Finally, because our assay is 3’-biased, we tested whether junction recovery decayed with distance from the 3’ end of the gene (Fig 8). Contrary to the naive expectation, recall did not fall toward the 5’ end. Among robustly expressed junctions (detected in at least 95 of the 190 bulk donors), recall of junctions within 500 bp of the 3’ terminus was 71.4%, slightly lower than the 80.2% for junctions 50 to 100 kb away, and repeating the analysis at transcript resolution, using the distance to the nearest containing transcript 3’ end (median 1,653 bp, versus 14,305 bp at the gene level), gave the same result (70.0% versus 81.4%). The gene-body coverage profile explained this position independence (Fig S6): across protein-coding genes 3 to 100 kb in length, single-cell coverage showed a sharp pileup at the 3’ end (a 6.2-fold excess in the 3’ relative to the 5’ decile) but was otherwise low and roughly flat across the gene body, so junctions along the length of expressed genes were sampled comparably. Taken together with the matched-bulk analysis, junction recovery in deep 3’ single-cell data is governed primarily by how strongly a junction is expressed and how many cells express it, not by its position within the transcript.

**Figure 8.**
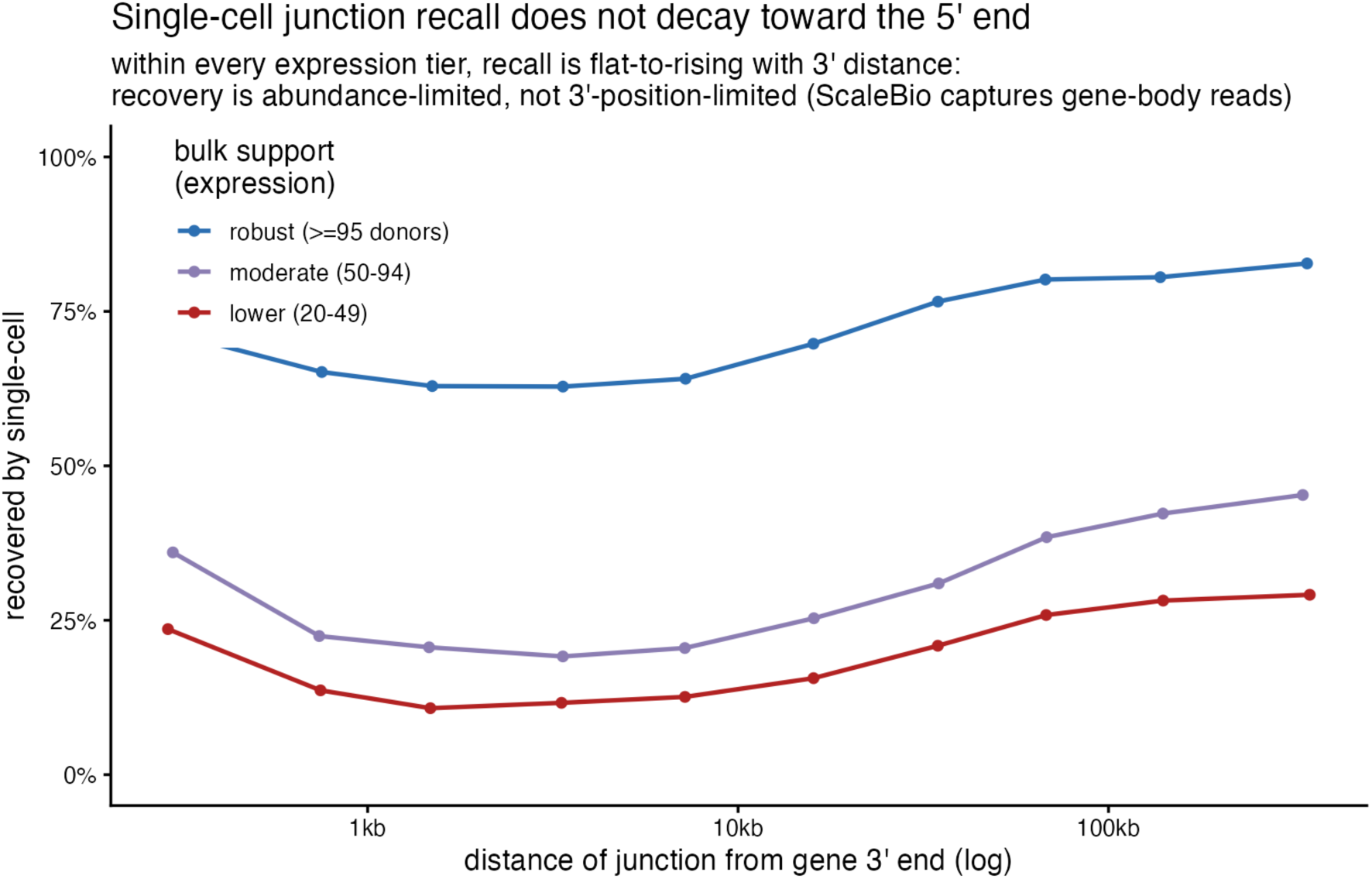
Junction recovery does not decay with distance from the 3’ end. Single-cell recall of bulk junctions in bins of distance from the gene 3’ terminus, stratified by bulk detection frequency. Recall did not fall toward the 5’ end (71.4% within 500 bp of the 3’ end versus 80.2% at 50 to 100 kb), consistent with a gene-body coverage profile that is flat apart from a sharp 3’ pileup.

These four axes together define a framework for designing splicing experiments in deep 3’ single-cell RNA sequencing: two design levers (depth targeting and budget allocation) and two constraints to plan around (the practical recovery ceiling and 3’-coverage bias). Figure 9 places them side by side with the measured design target for each.

**Figure 9.**
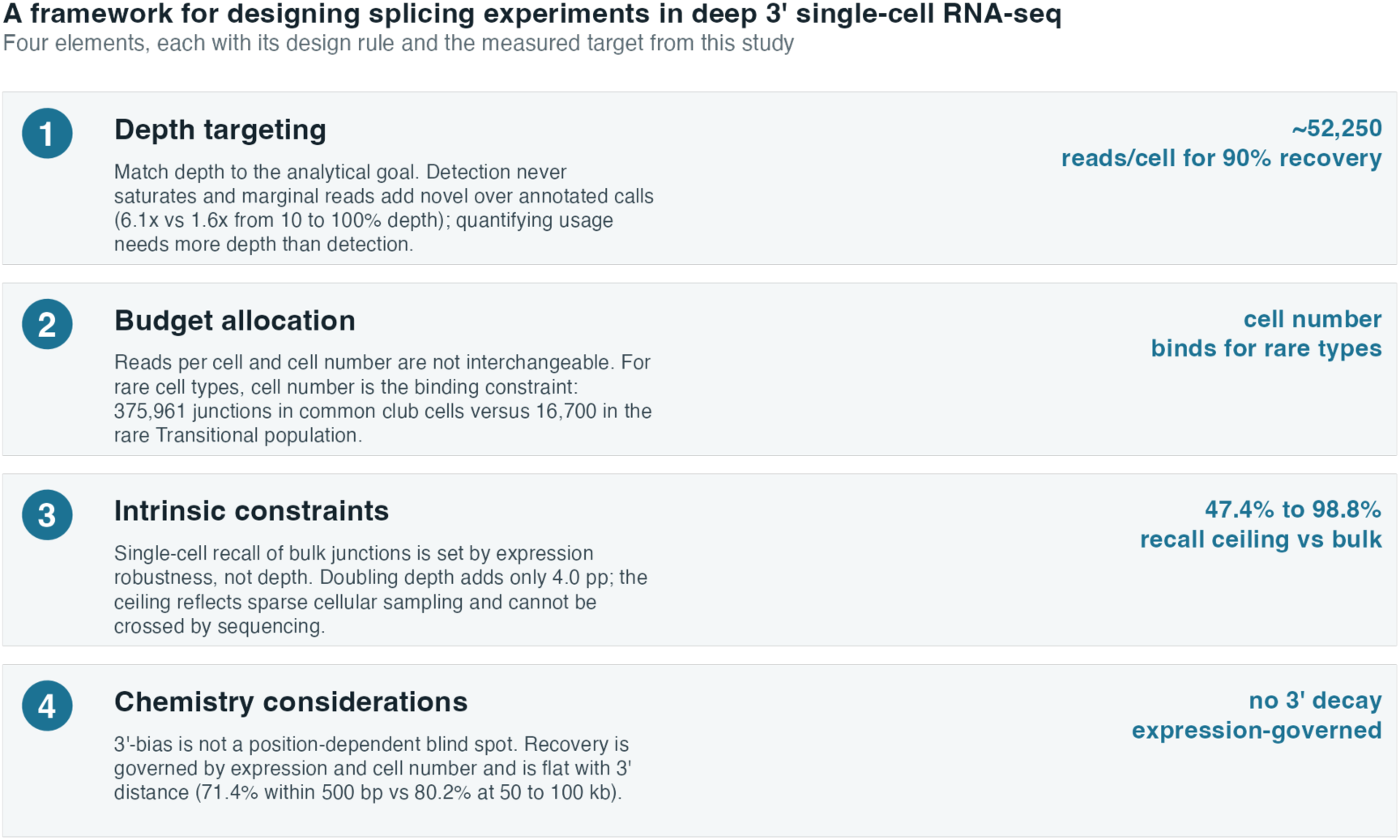
A framework for designing splicing experiments in deep 3’ single-cell RNA sequencing. Synthesis of the design framework. The four elements, each with its design rule and the measured target from this study: depth targeting (approximately 52,250 reads per cell for 90% recovery), budget allocation (cell number binds for rare types), practical constraints (a 47% to 99% recall ceiling against bulk), and chemistry considerations (no decay with 3’ distance).

### A worked example: what to expect at 50,000 reads per cell

To make the framework concrete, we applied it at a single realistic design point by downsampling the data to 50,000 reads per cell (near the depth of our own experiment) and asking how many matched-bulk junctions would be recovered in each cell type and at each expression level (Fig 10). At this depth, a common cell type of roughly 15,000 cells (club) recovered 158,786 bulk junctions, of which 65,742 were lowly expressed (fewer than ten reads per donor in bulk), whereas the rarest cell type of 68 cells (deuterosomal) recovered only 25,879 junctions, of which just 5,245 were lowly expressed. Recovery of highly expressed junctions (at least fifty reads per donor) was far more uniform across cell types (56,229 in club versus 14,267 in deuterosomal), so what rare populations lose is concentrated among the lowly expressed junctions. A reader planning an experiment can therefore read the expected junction recovery directly from the intended depth, the abundance of the target cell type, and the expression level of the junctions of interest.

**Figure 10.**
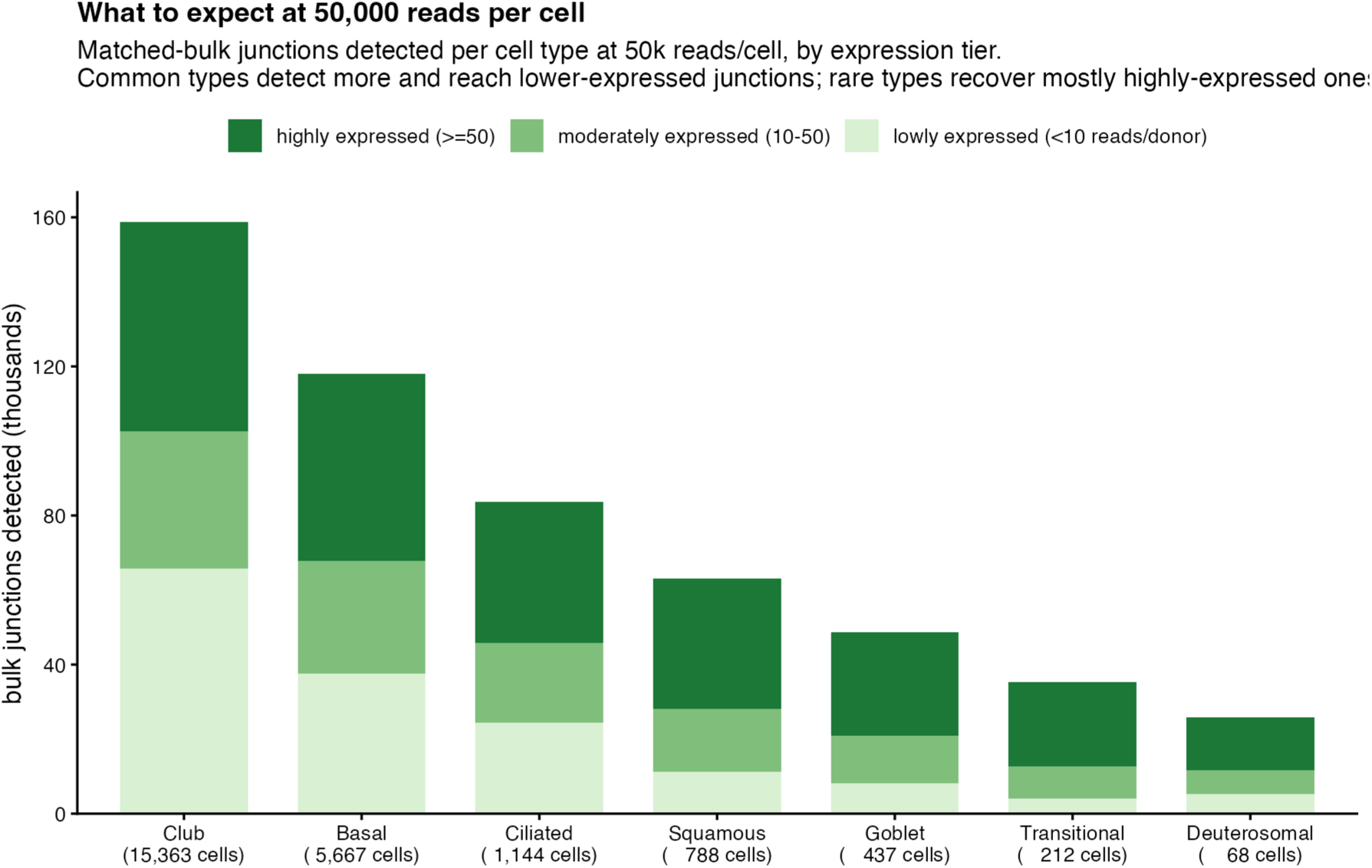
Worked example: junction recovery at 50,000 reads per cell. Matched-bulk junctions detected per cell type at 50,000 reads per cell, stacked by junction expression tier (highly expressed, at least 50 reads per donor; moderately, 10 to 50; lowly, fewer than 10 reads per donor). Cell types are ordered by abundance. Common cell types recover more junctions overall and reach lower-expressed junctions, whereas rare cell types recover predominantly highly expressed junctions.

### A sampling model predicts junction detectability

The detectability patterns of Figs 6 and 10 are captured by a simple sampling model, turning the framework into a predictive tool. Treating a junction as detected when at least one of its molecules is captured gives, under Poisson sampling, a detection probability that is a complementary-log-log function of the logarithms of three quantities: the junction’s expression (from bulk), the number of cells of the cell type, and the sequencing depth per cell. Fitting this model across the full grid of conditions yielded exponents of 0.81 for expression, 0.46 for cell number, and 0.49 for reads per cell, with similar fitted coefficients for cell number and reads per cell, quantitatively reinforcing the trade-off between the two design parameters. The model generalized across cell types: in leave-one-cell-type-out cross-validation, where the model predicts a cell type it never trained on, predicted and observed detection agreed with an R-squared of 0.94 and a mean absolute error of 1.3 percentage points (Fig 11). The fitted model serves as a calculator for experimental design: it predicts, for example, that a junction expressed at 100 reads per donor in bulk would be detected in 67% of cases in a common cell type of 5,000 cells sequenced to 50,000 reads per cell, but in only 23% of cases in a rare cell type of 200 cells at the same depth, and in 16% of cases for a junction ten times less expressed. The specific coefficients are calibrated to this tissue and chemistry, but the functional form provides a template that can be recalibrated for other systems.

**Figure 11.**
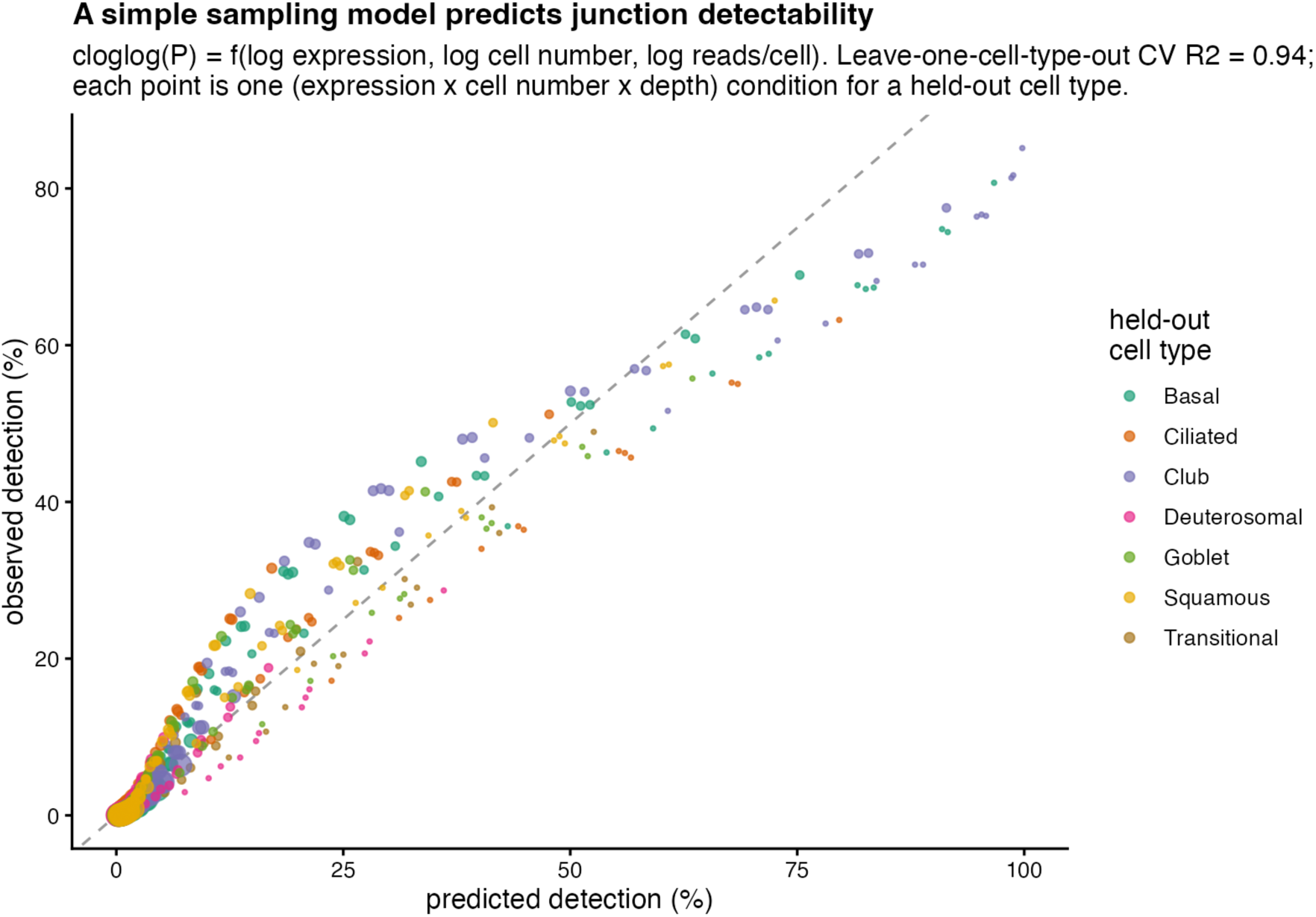
A sampling model predicts junction detectability. Observed versus predicted junction detection rate, with predictions from a complementary-log-log model of detection as a function of junction expression, cell number, and reads per cell. Each point is one (expression, cell number, depth) condition, colored by the held-out cell type in leave-one-cell-type-out cross-validation (each cell type predicted by a model trained only on the other cell types); the dashed line is the identity. Cross-validated R-squared is 0.94.

### Junction usage is cell-type-specific and concentrated in cytoskeletal genes

Finally, to move from junction detection to biology, we grouped junctions into intron clusters and tested for differential usage across the six cell types present in every donor, blocking on donor so that cell types were compared against biological replicates rather than pseudoreplicated cells. Of 12,209 testable alternative-splicing clusters, 40 clusters in 38 genes showed significant cell-type-specific junction usage (edgeR, FDR 0.05; Fig 12). The strongest signals fell in SPATS2L and the cytoskeletal linker DST, and in genes with well-established alternative splicing, including the tropomyosin gene TPM1 (FDR 9.3 x 10^-15) and the keratin gene KRT4, whose switch junction shifted by 75 percentage points in usage between cell types. Individual junctions tracked known cell identities, with FRMD6 usage specific to ciliated cells and SPINK5 to the secretory (club, goblet, and squamous) populations. At the pathway level, the gene sets ranked highest by nominal p value among the differentially spliced genes were dominated by cytoskeletal and cell-adhesion functions (actin binding, actomyosin, extracellular-matrix organization, and cell-substrate junctions; TPM1, TPM2, DST, SVIL, MYO6, ADD3, CD44), although no gene set survived genome-wide correction at this sample size. Cell-type-resolved single-cell data therefore recovers biologically coherent, cell-type-specific splicing that pseudobulk analysis of the same tissue would average away.

**Figure 12.**
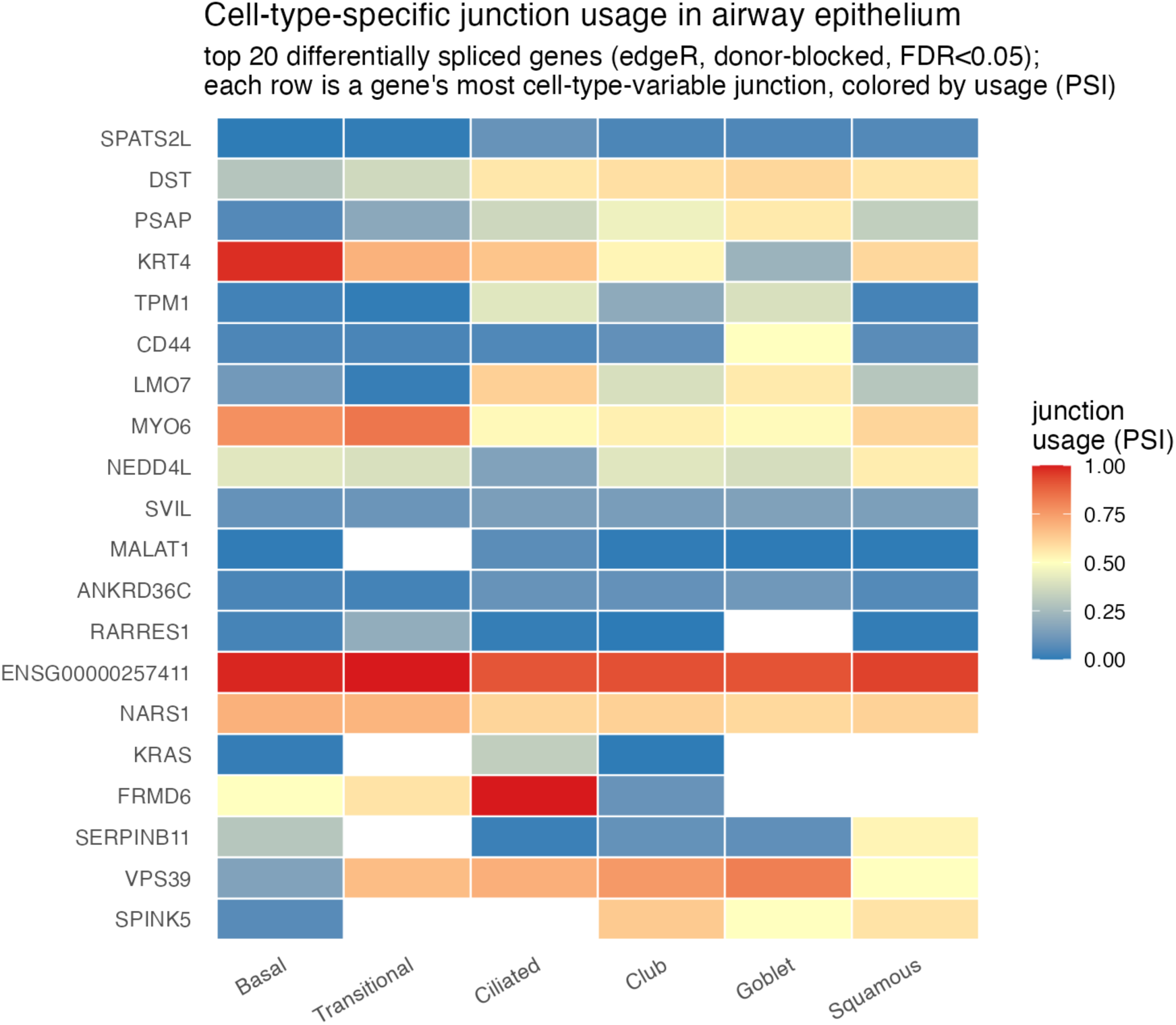
Cell-type-specific junction usage in the airway epithelium. Junction usage (percent spliced in, PSI) across the six cell types for the top 20 differentially spliced genes (edgeR, donor-blocked, FDR 0.05). Each row shows the most cell-type-variable junction of that gene’s intron cluster. Examples include KRT4 (basal), FRMD6 (ciliated), and SPINK5 (secretory).

### Two independent tools concur on the direction of cell-type splicing changes

To confirm that the cell-type splicing signal did not depend on a single analytic choice, we requantified the same junctions with MARVEL, a single-cell splicing tool that defines junction usage relative to total gene expression rather than within intron clusters. Comparing the two tools on the ciliated-versus-basal contrast, they agreed on the direction of change for 73% of the junctions that shifted appreciably (absolute PSI change of at least 0.05), rising to 80% among the junctions MARVEL called significant (Fig S7); the per-junction PSI changes were also positively but modestly correlated in magnitude (Spearman rho = 0.32 across 882 junctions quantified by both). The two quantifications are not numerically identical, because MARVEL normalizes junction counts to total gene expression whereas our intron-cluster approach normalizes within competing junctions, but they concur on the direction of cell-type splicing differences, indicating that the biological signal is robust to the quantification method.

## Discussion

Quantifying how sequencing depth governs splice-junction recovery turns a set of rules of thumb into measured design curves. Two features of the result were not obvious at the outset. First, there is no single depth at which junction detection is complete: total detection did not saturate even in our deepest samples, although highly expressed junctions in common cell types did approach saturation, so the appropriate target depends on the analytical goal and on the expression of the junctions of interest rather than on a universal threshold, and it is several-fold higher for quantifying splicing than for detecting it. Second, even ultra-deep sequencing of 12 samples did not recover all of the junctions seen in standard-depth bulk across 190 samples, a practical ceiling that reflects sample size and sparse cellular sampling rather than a shortfall that more sequencing would close. Together with the reads-versus-cells trade-off and the finding that 3’-coverage bias does not create a position-dependent blind spot, these results define a framework whose four elements are depth targeting, budget allocation, the practical recovery ceiling, and chemistry considerations.

### Practical guidance for experimental design

These findings translate into concrete recommendations. Detecting well-expressed junctions in common cell types is achievable at conventional depths, whereas comprehensive discovery of lower-expressed and previously unannotated junctions requires substantially deeper sequencing, and quantifying splicing usage requires close to full depth; studies quantifying splicing at reduced depth should restrict to well-covered clusters or aggregate across cells and samples. For rare cell types, depth cannot substitute for cell number, so designs targeting rare-cell splicing should prioritize cell enrichment through sorting, spatial selection, or higher-throughput profiling. The returns from additional sequencing diminish steeply near the ceiling (Fig S8), and because the practical ceiling against bulk reflects sparse cellular sampling rather than depth, the lowly and variably expressed junctions that single-cell data miss are more effectively recovered by adding cells than by adding reads once sequencing is already deep, so studies aiming to catalog rare splicing events will need to combine single-cell resolution with either targeted enrichment or a matched bulk reference. Our empirical estimate of approximately 52,250 reads per cell fills the design regime between conventional gene-level profiling (∼15,000 reads/cell [12]) and full-length isoform sequencing in Smart-seq (1 to 4 million reads/cell [16]), and the sampling model presented above (Fig 11) provides a calculator that can be applied to a planned experiment or recalibrated for other systems.

### Cell-type-specific splicing biology

Beyond the methodological contributions, our cell-type-resolved analysis identified 40 significantly differentially spliced intron clusters concentrated in genes with cytoskeletal and cell-adhesion functions, consistent with the known role of alternative splicing in epithelial cell identity and remodeling. These significant clusters were strongly enriched among highly expressed genes (median expression at the 93rd percentile of tested clusters, roughly elevenfold above clusters that did not reach significance), so the analysis is powered mainly to detect cell-type-specific splicing in abundant genes and lower-expression programs likely remain undetected. The modest hit count also reflects the current sample size (five donors, six cell types with matched biological replicates); comprehensive cataloging of cell-type-specific splicing in the airway epithelium will require larger cohorts.

### Generalizability and limitations

The framework’s qualitative principles should generalize broadly. Because junction detection is fundamentally a sampling problem, the functional form of the detectability model (detection increasing with expression, cell number, and reads per cell with diminishing returns), the reads-versus-cells trade-off, the observation that rare cell types are cell-number-bound rather than depth-bound, and the existence of a practical recovery ceiling against bulk are direct consequences of sparse sampling and should generalize across single-cell platforms and tissues. The specific quantitative targets we report, however, are calibrated to the ScaleBio combinatorial chemistry and the airway epithelium: the 52,250 reads per cell required for 90% recovery, the 47.4% recall ceiling, the model coefficients (∼0.8 for expression, ∼0.5 for cells and reads), and the 3’-distance independence are chemistry- and tissue-specific and will need re-estimation for other droplet platforms (10x 3’, 10x Flex), for tissues with different cell-type complexity, and especially for chemistries with substantially different coverage profiles (for example, full-length Smart-seq3 or 5’-based methods, where the 3’-distance-independence we observe should not be expected to hold). The complementary-log-log functional form of the sampling model provides a template for such recalibration. Our study has additional limitations. The matched bulk reference was derived from basal airway epithelial cells cultured in monolayer rather than from the differentiated cultures used for single-cell profiling; because basal cells in monolayer may differ biologically from basal cells within a differentiated epithelium, this difference likely contributes to the single-cell-versus-bulk PSI gap, although the within-donor comparison of single-cell samples against the same donor’s bulk reproduced the recall ceiling almost exactly, arguing against a substantial confounding of the recovery estimates. Depth downsampling was performed within samples rather than across a controlled empirical depth ladder, so cross-sample comparisons are confounded by biology and technical variation. And the cell-type-specific splicing analysis is powered to detect only the most prominent programs.

### Conclusions

Single-cell RNA sequencing offers a resolution advantage for splicing analysis that bulk methods cannot match, but this resolution comes with sampling costs that fundamentally constrain what can be detected. By quantifying these constraints in a deeply sequenced 3’-biased single-cell dataset, we provide the empirical foundation for designing single-cell splicing experiments and for interpreting their results. Studies planning depth targets, choosing between more reads or more cells, or evaluating the completeness of junction recovery can now do so against measured design curves rather than heuristic estimates. The deep single-cell airway epithelial dataset with matched bulk that we release as part of this work provides a resource for benchmarking future single-cell splicing methods against ground truth.

## Author Contributions

A.S. conceived and designed the study, performed data and statistical analyses, acquired funding, and wrote and revised the manuscript. C.L., A.S.V., and K.B. contributed to data collection. J.C. contributed to data collection and data and statistical analyses. Y.T. contributed to study conception and design, data collection, and funding acquisition. P.J.C. contributed to study conception and design, funding acquisition, and manuscript writing and revision. All authors reviewed and approved the final manuscript.

